# Spontaneous oscillatory activity in episodic timing: an EEG replication study and its limitations

**DOI:** 10.1101/2025.09.03.674038

**Authors:** Raphaël Bordas, Virginie van Wassenhove

## Abstract

Episodic timing refers to the one-shot, automatic encoding of temporal information in the brain, in the absence of attention to time. A previous magnetoencephalography (MEG) study showed that the relative burst time of spontaneous alpha oscillations (α) during quiet wakefulness was a selective predictor of retrospective duration estimation. This observation was interpreted as α embodying the “ticks” of an internal contextual clock. Herein, we replicate and extend these findings using electroencephalography (EEG), assess robustness to time-on-task effects, and test the generalizability in virtual reality (VR) environments. In three EEG experiments, 147 participants underwent 4-minute eyes-open resting-state recordings followed by an unexpected retrospective duration estimation task. Experiment 1 tested participants before any tasks, Experiment 2 after 90 minutes of timing tasks, and Experiment 3 in VR environments of different sizes. We successfully replicated the original MEG findings in Experiment 1 but did not in Experiment 2. We explain the lack of replication through time-on-task effects (changes in α power and topography) and contextual changes yielding a cognitive strategy based on temporal expectation (supported by a fast passage-of-time). In Experiment 3, we did not find the expected duration underestimation in VR, and did not replicate the correlation between α bursts and retrospective time estimates. Overall, EEG captures the α burst marker of episodic timing, its reliability depends critically on experimental context. Our findings highlight the importance of controlling experimental context when using α bursts as a neural marker of episodic timing.

**Significance Statement:** How does the brain automatically keep track of time during everyday experiences? This study investigates alpha brain activity as a marker of contextual changes during quiet wakefulness. We successfully replicated the original findings using EEG, which is more widespread than MEG, but found some limitations. This neural marker is sensitive to mental fatigue and experimental context, with participants adopting temporal expectation strategies that alter the relation between alpha and temporal estimation. Virtual reality environments also affected behavior in a way that suggested prospective timing which the marker is known not to capture. As alterations in timing affect numerous neurological and psychiatric conditions, establishing a robust neural marker of experiential time has important implications for both basic neuroscience and clinical applications.

## Introduction

Most of our temporal experiences are lived as one-shot moments or episodic experiences of events. The term “episodic timing” was recently used to refer to the automatic encoding of temporal information in the absence of task requirements or overt attention to time (Azizi et al., 2023). Episodic timing focuses on the neural mechanisms encoding the temporal relations between events (e.g., duration, order, speed) ultimately stored in, and constitutive of, the *when* of memory. Neural indices of episodic timing should be indicative of participants’ subsequent duration memory, which can be probed through a retrospective estimation of elapsed time. In a recent study, participants were recorded with magnetoencephalography (MEG) during quiet wakefulness for a few minutes after which they were unexpectedly asked to report as precisely as possible how long the recording episode was. The authors reported that the increase in total alpha power (α, 8 - 12 Hz) during resting-state was linearly predictive of participants’ retrospective time estimate. More specifically, the relative α burst time was the best predictor of retrospective duration estimation (Azizi et al., 2023). This relation vanished during prospective timing (when people knew they would have to report the duration of the recording, thus paying attention to time). α burst time was postulated to be a neural index of episodic timing.

Retrospective duration tasks are single-shot duration estimations (Balcı et al., 2023; Chaumon et al., 2022), which have been postulated to engage episodic memory processes (Block, 1985; Hicks et al., 1976; Michon, 1975). The contextual change hypothesis (Block, 2014; MacDonald, 2014; Ornstein, 1969) posits that retrospective duration estimates depend on the amount of contextual change participants experienced. Spontaneous α bursts reflect discrete shifts in brain states, which could serve as neural markers of such internal contextual changes during quiet wakefulness (Azizi et al., 2023). The fact that the relation between relative α burst time and duration was only observed retrospectively and not prospectively aligned well with this hypothesis according to which retrospective duration is reconstructed from memory, not tracked in real-time. The authors suggested that spontaneous α bursts embody the “ticks” of an internal contextual clock, which operationalized the concept of contextual change at the neural level (Azizi et al., 2023).

α oscillations (Berger, 1929) occur spontaneously (Buzsáki & Draguhn, 2004) and their functional role has been refined over time (Bazanova & Vernon, 2014). The notion of cortical idling (Pfurtscheller et al., 1996) was proposed following the seminal observation that closing the eyes increased posterior α (Adrian & Matthews, 1934). It evolved towards the functional inhibition hypothesis as a regulatory mechanism of sensory processing (Jensen et al., 2012; Jensen & Mazaheri, 2010; Klimesch et al., 2007; von Stein et al., 2000). In quiet wakefulness, fluctuations of α activity capture moment-to-moment changes of brain activity in response to external stimulations but also to internally-driven cognitive processes (Halgren et al., 2019; Sadaghiani & Kleinschmidt, 2016). In internally directed thoughts such as in mind-wandering, participants are oriented towards their internal stream of thoughts rather than to external stimuli (Smallwood & Schooler, 2015). Accordingly, the dynamic changes in α fluctuations co-occur with periods of mind-wandering: for instance, α power preceding a probe is higher when participants mind-wander than when they do not (Compton et al., 2019). Mind-wandering can be interpreted as a lapse of attention, which yields to an underestimation of the duration of sensory stimuli (Terhune et al., 2017).

Herein, we wished to extend the original MEG observations. First, we wished to replicate and generalize the protocol with EEG, a widely used technique that can easily adapt to ecological situations (Stangl et al., 2023; Vallet & van Wassenhove, 2023) and clinical settings. This is important as this episodic timing marker can serve clinical diagnosis seeing that time perception is altered in numerous pathologies (Hinault et al., 2023). Second, we assessed the robustness of the episodic timing marker to time-on-task effects, which are characterized by a large-scale increase in α power and/or a decrease in peak frequency (Benwell et al., 2019). Third, we tested the replicability in Virtual Reality (VR), increasingly used in time perception research (Jording et al., 2022; Lamprou-Kokolaki et al., 2024; Tobin et al., 2010; Tobin & Grondin, 2009).

In all EEG experiments, participants were recorded with open eyes in quiet wakefulness for 4 minutes in the absence of any task requirement; at the end of the recording, participants reported as precisely as possible the duration (in minutes, seconds) of the recording that just occurred (Azizi et al., 2023). Participants were recorded before any task (Exp. 1), after performing timing tasks for 90 minutes (Exp. 2) or in VR before any task (Exp. 3). We replicate the original findings when participants perform the retrospective timing task before any other task, but not after an experimental session nor in VR.

## Materials and Methods

### Participants

A total of 147 participants were recruited to take part in the study (71 males, age = 25 years old, +/− 5 years). All participants had normal or corrected-to-normal vision, were right-handed, and were naive as to the purpose of the study. None had known neurological or psychiatric disorders, and none were under medical treatment. All participants provided a written informed consent form validated by the Ethics Committee for Research (CER) of Paris-Saclay University (CER-Paris-Saclay-2018-034-A2 and CER-Paris-Saclay-2023-089) or the Research Ethics Committee of Neurospin, CEA (CPP NeuroSpin CEA 100 049 2018).

31 participants took part in Experiment 1 (Exp. 1) and performed the retrospective timing task *before* taking part in other tasks. Two participants were excluded due to inconsistent behavior, and two were excluded because they used a counting strategy against our instructions (see below). One participant was excluded because the scalp topography was outside the group distribution (see Supp Fig. 1). Thus, 26 participants (15 males, age = 27 years old, +/− 5 years) were included in the final analysis of Exp. 1.

In Experiment 2 (Exp. 2), 51 participants performed the retrospective timing task ∼90 minutes after performing an explicit or an implicit timing task. Two participants were excluded due to technical issues in the EEG recordings, and one participant was excluded because of a counting strategy. Three additional participants were excluded because their topography was outside the group distribution (see Supp Fig. 1), leaving 45 participants analyzed in Exp. 2 (20 males, age = 23 years old, +/− 5 years). Among these 45 participants, 21 were recorded during a resting-state alternating eyes-closed and eyes-open before taking part in an explicit timing task. This subset of data was analyzed to compare the α burst before and after an explicit timing task (analysis reported in Fig. 5).

In Experiment 3 (Exp. 3), 65 participants performed the retrospective timing task in a VRl environment before taking part in subsequent tasks. Before doing so, and following a short VR practice session, participants filled out the Simulator Sickness Questionnaire (Bouchard et al., 2007; Kennedy et al., 1993) to ensure they were fit for the study. One participant was excluded for health reasons, two for missing data, one for bad EEG data, and one because the scalp topography was outside the group distribution. Three additional participants were excluded because they were outliers in the behavioral data distribution (see Methods). Thus, a total of 57 participants (25 males, age = 25 years old, +/− 4 years) were effectively analyzed in Exp. 3.

The required sample size was computed using an *a priori* power analysis of Pearson’s correlation coefficients based on a published study (Azizi et al., 2023). We aimed for a power greater than 0.8, leading to a minimal sample size of 23 participants when considering a positive effect of rho = 0.5 (one-sided test). The number of participants recruited in each experiment was well above the needed sample size indicated by the power analysis.

In total, 132 participants were included in the final analysis in this study (60 males, age = 25 years old, +/− 5 years).

### Experimental design

The retrospective time task consisted of an eye-open resting-state EEG recording followed by a question and a questionnaire.

In Exp. 1 and Exp. 2, participants were seated in front of a distant wall. They were asked to stare in front of them at a point on the wall so as to attenuate eye movements, to stay as still as possible, to relax their body, and to avoid muscle and jaw tensions. They were asked to keep their eyes open and to “clear their minds”. At the end of the resting-state recording, participants were unexpectedly asked: “How much time has elapsed since the beginning of the recording? Be as precise as possible in minutes and seconds”. They were asked to report the strategy they used to make their duration estimation. The experimenter inquired participants as to whether they were paying attention to time during the resting-state or whether they guessed they would be asked to estimate the duration of the recording. None of the participants reported having done so.

In Exp. 1, participants were recorded for 4 or 5 minutes and performed the retrospective task before any other experimental task. In Exp. 2, participants underwent a 4-minute resting-state EEG recording after completing a 90-minute timing experiment. The timing experiment could involve explicit attentional orientation to time using a visual temporal adaptation task (e.g., Jin et al. (2024)) or be an implicit timing task (e.g., Herbst et al. (2022)).

A subset of participants in Exp. 2 (N = 21) was recorded with a 3-minute resting-state recording, alternating eyes open and eyes closed every 30 seconds before the actual explicit timing experiment. No duration estimation was asked of participants for this recording, and no indication of timing was provided. We exploited the 90-second eyes-open data for a control analysis reported in Fig. 5.

In Exp. 3, participants were tested while seated in either a small or large virtual environment (Fig. 2A). Each resting-state started with a short exploration phase of the virtual environment (i.e., 20 seconds). Then, participants were instructed to stay as still as possible, to clear their minds, and to stare at the lamp positioned on a table in front of them to avoid any eye movements. The start and end of the recording were indicated to the participant by the lamp, which lit up green twice: once to indicate the start of the recording, a second time to indicate the end of the recording. At the end, participants removed the VR headset and were asked the same question as in Exp. 1 and Exp. 2 and to fill out the same questionnaire as in Exp. 1 and Exp. 2.

### EEG acquisition

49 EEG datasets (26 in Exp. 1; 23 in Exp. 2) were acquired using a 32-channel EasyCap system (Brain Products, GmbH) at a sampling frequency of 500 Hz, in DC (no high-pass filter) with a 131 Hz low-pass filter applied online. Impedance was kept below 25kΩ. 79 EEG datasets in Exp. 2 and Exp. 3 were acquired with an eego mylab amplifier and a 64-channel Waveguard cap (ANT Neuro, Enschede, Netherlands) at a sampling frequency of 1 kHz. EEG recordings were collected in DC (no high-pass filter) with an online low-pass filter of 260 Hz. The reference electrode was CPz. The vertical electrooculogram (EOG) was acquired with one electrode positioned above the left eye. Impedance was kept below 20kΩ.

### EEG preprocessing

Data from all participants were identically preprocessed with a custom pipeline based on MNE Python (Gramfort et al., 2013). To follow the procedure of the previous study (Azizi et al., 2023), and unless mentioned otherwise, the first 4 minutes of every recording were considered for analysis to homogenize the duration of the recordings across all analyses.

First, bad channels were manually marked upon visual inspection of the continuous raw signals. Mastoids were noisy and did not contain relevant brain activity (Cohen, 2014); we excluded them from further analysis. Second, we band-pass filtered all channels between 1 Hz and 130 Hz with a notch filter set at 50 Hz and 100 Hz to remove the power line noise and its harmonics. Third, bad channels were interpolated. Finally, data were re-referenced to the average of all channels.

To remove ocular artifacts, one copy of the signals was high-pass filtered at 1.5 Hz, downsampled to 500 Hz, and passed on to a FastICA algorithm (Gramfort et al., 2013). The classification of components containing EOG artifacts relied on a Pearson correlation between the filtered data and the filtered vertical EOG channel placed above the left eye. The automatic classification of EOG artifacts was double-checked by visual inspection through all components and manually corrected when necessary. The ICA with all the bad independent components removed was then applied to the continuous signal filtered between 1 Hz and 130 Hz. The median number of components removed by participants was 2, with a maximum of 7.

To further ensure the quality of our data, data containing muscular artifacts were marked and rejected automatically using native MNE functions. A muscular artifact was detected when the sum of envelope magnitude across all channels divided by the square root of the number of channels in a segment exceeded a threshold set to 10 for data recorded with the EasyCap system and 15 for data recorded with the eego mylab system. To prevent a high number of small data cropping and edge artifacts in burst detection, the minimum length of a good data span was set to 500 ms, and so was the minimum length of a bad data span. The remaining artifacts were marked manually and rejected during the final visual inspection. This led to up to 5 seconds rejected per recording in Exp. 1 (M = 1.70, SD = 1.64), 8 seconds in Exp. 2 (M = 1.97, SD = 2.61), and up to 28 seconds rejected in Exp. 3 (M = 4.73, SD = 5.97). Critically, to take this into account, all temporal EEG measures were relative to the recording duration after preprocessing.

### Merging datasets acquired with different systems

All recordings acquired at a sampling rate higher than 500 Hz were downsampled at 500 Hz to ensure a homogenous sample count across analyses. The intersection between montages of 64 and 32 channels was retained for all analyses, thus including 27 channels from the 10-10 international system.

### Spontaneous oscillatory spectrum and localizers

To detect α activity (*8 Hz – 12 Hz*) at every channel, we used a custom Python implementation of the IRASA algorithm (Wen & Liu, 2016). IRASA is used to separate the oscillatory component from the aperiodic one (1/f slope), typically seen in the power spectral density (PSD) of brain activity (e.g., Fig. 1A). First, the continuous EEG signals were divided into fixed-length epochs of 5 seconds to allow a 0.2 Hz frequency resolution in the spectral domain. Second, the IRASA algorithm up-sampled and down-sampled the epoched data at pairwise non-integer scaling factors *h* and *1/h*. Specifically, the sampling frequency was respectively multiplied by *h* and *1/h* with *h* ranging from 1.1 to 2.95 in increments of 0.05. Third, for each (*h*, *1/h*) pair, the PSD was computed using Welch’s periodogram between 1 Hz and 45 Hz and the IRASA algorithm took the geometric mean of these two PSDs. Fourth, the aperiodic component was estimated from the median of all resulting PSDs. As a result, we obtained two estimations: one for the aperiodic component and one for the oscillatory component. The oscillatory spectrum resulted from the subtraction of the aperiodic component from the original PSD. The α peak power was defined as the maximum amplitude of the oscillatory spectrum between 8 Hz and 12 Hz. The individual α peak frequency (iAPF) was defined as the argument of this maximum. The power and the iAPF of α activity were used for sanity checks in Exp. 2.

**Figure 1.**
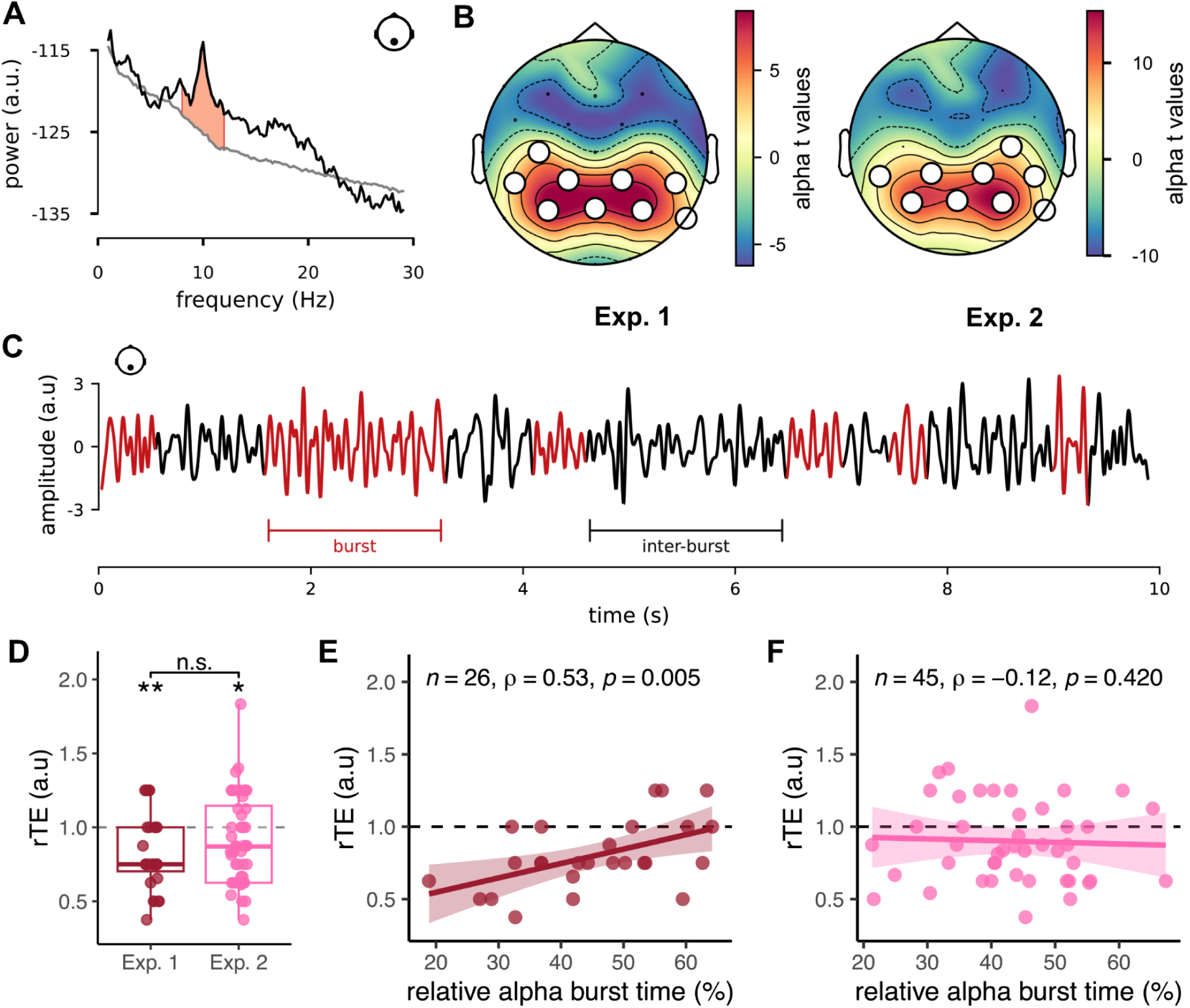
α burst time correlates positively with retrospective time estimates (rTEs) when recorded before (Exp. 1), but not after (Exp. 2), a timing task. **(A)** Spectrum decomposition using IRASA (Wen & Liu, 2016). The black line is the original spectrum computed from a single parietal channel (Pz). The grey line illustrates its aperiodic component. The red-shaded area represents the integrated power of the oscillatory spectrum between 8 Hz and 12 Hz (α activity). **(B)** Localization of α activity in the spectral domain during a resting-state acquired before (Exp. 1) or after a timing task (Exp. 2). Topographic maps in both experiments show the spatial distribution of the t-values obtained when testing the integrated α power at each channel against the average of all channels. The white dots denote the cluster retained in all subsequent analyses and experiments. Topographies of participants that were excluded based on this localization can be found in Extended Data Figure 1-1. **(C)** Example of α bursts recorded at Pz during resting-state. A burst is defined as a series of contiguous cycles with a consistent period, amplitude, and monotonicity in the α range. Each burst is shown in red. The relative burst time is the percentage of cycles classified as belonging to a burst in the entire time series. **(D)** Distribution of the relative time estimates before any task (N = 26; Exp. 1; red) and after performing a 90-minute task (N = 45; Exp. 2; pink). Each dot is a participant (single-trial experiments). Standalone stars indicate a significant temporal underestimation according to a Wilcoxon rank-sum test (**p = 0.002; *p = 0.028) with the red dotted line as the null hypothesis value (i.e. estimated duration equals veridical duration). n.s. indicates the absence of a significant difference between Exp. 1 and Exp. 2 (Wilcoxon rank-sum test, W = 545.50, p = 0.257). The homogeneity of participants grouped in Exp. 2 is assessed in Extended Data Figure 1-2. The PoTJs associated with these duration estimates are shown in Extended Data Figure 1-3. **(E-F)** Spearman’s correlation between the relative α burst times and the rTEs in Exp. 1 (E) and Exp. 2 (F). The straight line is the regression line, and the shaded area is the 95% CI. Sensitivity analysis of the linear model of panel (C) can be found in Extended Data Figure 1-4.

To control for broadband frequency variations, we used the slope of the aperiodic component. It was obtained via a linear regression in the log-log space, following methods from (Donoghue et al., 2020). In that space, the aperiodic component 𝑦_𝑎𝑝_ is assumed to be expressed as 𝑦_𝑎𝑝_ = 𝑏 − χ𝑙𝑜𝑔(𝐹) with 𝐹 the frequency, χ the slope, and 𝑏 the intercept. In our analysis, only models with R^2^ > 0.9 were retained to ensure the robustness of the statistics.

Once the oscillatory spectra were obtained, we identified the most sensitive channels to the α activity. For this, a cluster-based permutation t-test (Maris & Oostenveld, 2007) was performed on the integrated power of the oscillatory spectrum in the α range (Fig. 1A). Specifically, we tested whether each EEG channel had an integrated power higher than the average of all EEG channels using t-statistics. The null distribution of this t-test was obtained through a Monte-Carlo approximation with N = 4000 permutations. Using the channels connectivity matrix estimated with the Delauney triangulation implemented in MNE-Python (Gramfort et al., 2013), clusters of interest were defined as neighboring channels whose upper-tail t-statistics exceeded a threshold *q^(N−1)^*. This threshold was set at the 99.9^th^ percentile of a Student’s t-distribution with degrees of freedom N - 1, with N the number of participants. For Exp. 1, 𝑞^(26)^ = 2. 48, for Exp. 2, ^(47)^ = 2. 41, and for Exp. 3 ^(57)^ = 2. 39. The p-value was estimated based on the proportion of permutations exceeding the observed maximum cluster-level test statistic. Only clusters with corrected p-values < 0.05 are reported. All clusters were located in the parietal channels (Fig. 1B for Exp. 1 and 2, and Fig. 2B for Exp. 3) and are referred to hereafter as “parietal clusters”. One participant in Exp. 1, three participants in Exp. 2, and one participant in Exp. 3 displayed no oscillatory activity at the group-level cluster. Hence, they were excluded from further analysis (see their profiles in Extended Data Fig. 1-1). Thus, as stated in Participants, N = 26 in Exp. 1, N = 45 in Exp. 2, and N = 57 in Exp. 3 were included in our final analyses (see Table 1).

**Figure 2.**
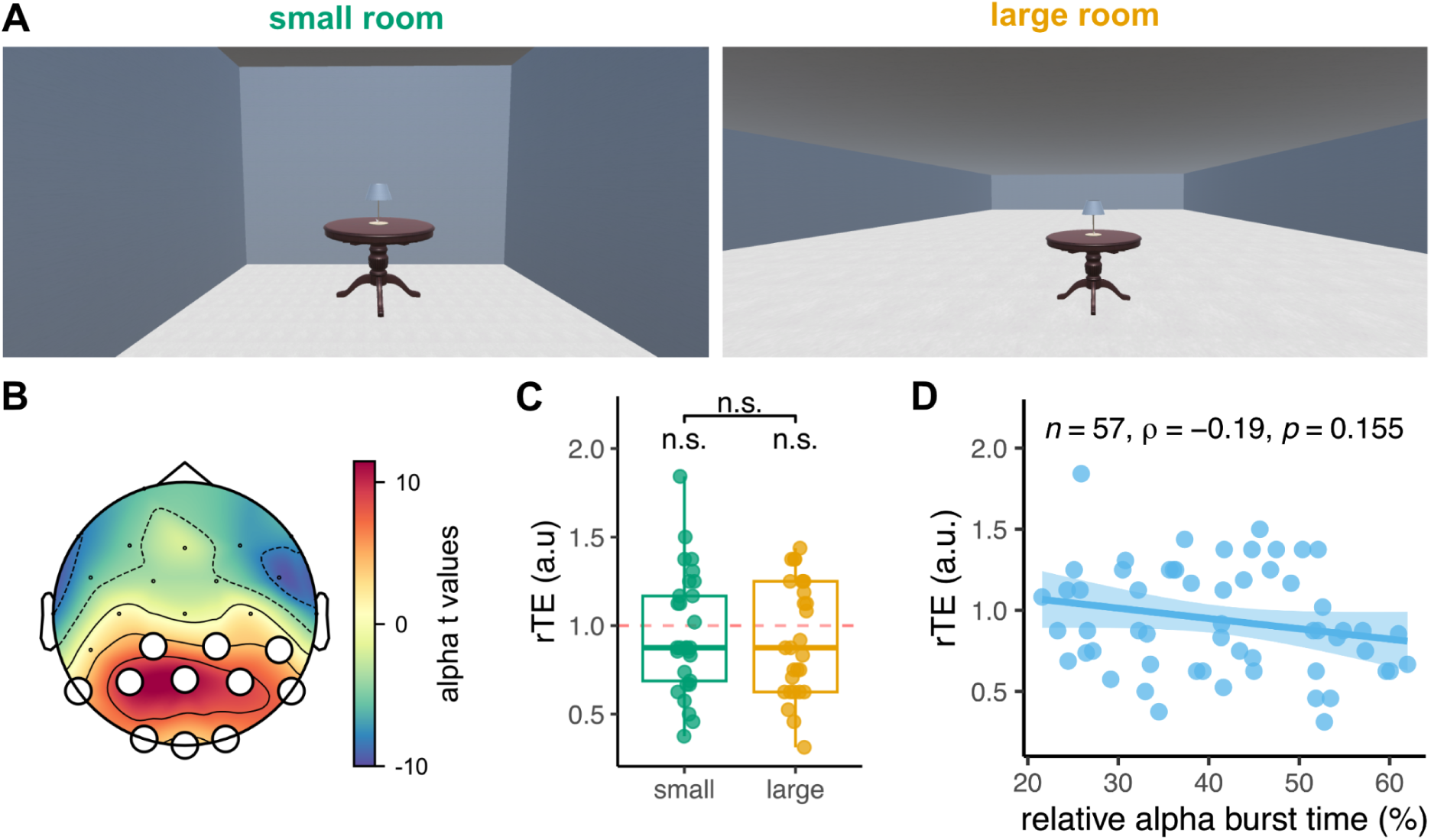
α burstiness does not predict retrospective duration in VR (Exp. 3). **(A)** Participants were seated for 4 minutes in a small room (N=29; left panel) or a large room (N=28; right panel). The lamp turned green for 500 ms at the beginning and at the end of the interval. **(B)** Localization of α activity in the spectral domain. The topographic map shows the distribution of the t-values obtained when testing the integrated α power at each channel against the average of all channels. The white dots denote the cluster retained in all subsequent analyses. **(C)** Distribution of the rTEs in the small (N = 29; green) and large (N = 28; yellow) rooms. Each dot is a participant. No significant underestimation was found in either condition (standalone n.s. markers). No significant differences were found between experimental conditions, as assessed by a two-sample unpaired Wilcoxon test. **(D)** Pearson’s correlation between the relative α burst times and the rTEs. The straight line is the regression line, and the shaded area is the 95% CI.

**Table 1.** Sample size in each experiment and for each analysis after the exclusion of participants who did not display the frequency band of interest according to the localizer analysis.

| | $\alpha$ (8 - 12 Hz) | $\theta$ (3 - 7 Hz) | $\beta$ (15 - 30 Hz) |
| --- | --- | --- | --- |
| Exp. 1 | 26 | 24 | 23 |
| Exp. 2 | 45 | 44 | 44 |
| Exp. 3 | 57 | 56 | 54 |

The same procedure was separately repeated for θ (*3 Hz – 7 Hz*) and β (*15 Hz – 30 Hz*) activity. Table 1 summarises the resulting sample size for each frequency band.

### Cycle-by-cycle analysis and burst detection

To compute the bursting features of continuous signals, we used the bycycle toolbox (Cole & Voytek, 2019). The bycycle algorithm computes several statistics on each cycle of a periodic time series to identify burst episodes. The settings for the detection algorithm used identical values as a previous study (Azizi et al., 2023): the amplitude fraction threshold was set to 0.2; the amplitude consistency threshold to 0.4; the period consistency threshold to 0.4, the monotonicity threshold to 0.8, and the minimum number of cycles to 3.

For the α bursts, and prior to the thresholding algorithm, the signal was band-pass filtered between 4 Hz and 16 Hz. The bandwidth larger than the 8 Hz - 12 Hz definition of α avoided strict transition bands (border artifacts). Outcomes of the bycycle algorithm were modified as follows: burst cycles with a frequency not included in (8 Hz - 12 Hz) were discarded, unless they were surrounded by two burst cycles and the mean period of the burst was included in the extended α range (7.5 Hz - 12.5 Hz). This procedure avoided splitting bursts due to the period variability from one cycle to another inside a burst. The outcome of this procedure was the relative burst time, which corresponds to the ratio of the number of cycles that are part of an α burst to the total number of cycles in the signal (Fig. 1C). As a possible interpretation of this variable, the more bursty the signal is, the more stationary the signal is in the time domain. On the contrary, a 0% bursting signal means that no α activity is detected.

Additionally, to assess the stability of α burstiness over time, the process described above was repeated on nine 25-second non-overlapping time windows. This window duration took into account the muscular artifact rejection that shortened some recordings by a few seconds and ensured identical signal length for all participants (see EEG processing paragraph).

We used the same procedure to detect bursts in the θ and β bands, with adapted thresholds and parameters. For θ (3 Hz - 7 Hz), the monotonicity threshold was set to 0.7, the amplitude fraction threshold to 0.3, the minimum number of cycles to 2, and signals were pre-filtered between 2 Hz and 8 Hz. For β (15 Hz - 30 Hz), the monotonicity threshold was set to 0.9, and both period and amplitude consistency thresholds to 0.5.

### Statistical analysis

All statistical analyses were performed using the R Software version 4.3.1 (R Core Team, 2021).

#### Behavioral data

Relative time estimates (rTEs) were computed as the ratio between duration estimation and clock time. Three outliers in Exp. 3 were detected when defined as values outside the interval [− 1. 5×𝐼𝑄𝑅, 1. 5×𝐼𝑄𝑅], where 𝐼𝑄𝑅 is the interquartile range. No outliers were detected in Exp. 1 or Exp. 2. Because the rTEs were not normally distributed (as assessed by visual inspection of the Q-Q plots), the hypothesized underestimation of retrospective duration was assessed using a Wilcoxon signed-rank test. Likewise, rTE differences between experiments (Exp. 1 *vs.* Exp. 2) and experimental conditions (small *vs*. large room in Exp. 3) were analyzed using a Wilcoxon rank-sum test.

#### Regressors

Due to the non-normal distribution of the rTEs, correlations were performed using Spearman’s coefficient (denoted ρ). The corresponding linear model was also computed to assess the robustness of the relation between each predictor and the dependent variable. Tested predictors were the relative burst time, the mean period of burst cycles, and the slope of the aperiodic component. Then, Cook’s distance D was calculated to identify influential observations. Participants with D > 4/N were marked as potential outliers, with N the total number of observations. The model was rerun without these observations to ensure no individual points drove the findings.

Ordinal logistic regressions were used for the models testing passage of time judgments using the PoTJ as the dependent variable and the experiment (Exp. 1, Exp. 2, or Exp. 3) as the regressor (see Extended-Data Figure 1-3). The significance of ordinal regressions was assessed through the likelihood ratio test against the null model.

#### Sensitivity analysis

Starting with 5 participants up to 26 (our lowest achieved sample in Exp. 2) and by steps of 1, we computed the linear model 𝑟𝑇𝐸 = 𝑏_0_ + 𝑏_1_𝑥 (with 𝑥 the burst time) using randomly sampled data without replacement. We repeated the process 100 times to get an average t value testing the coefficient 𝑏_1_ against zero for each sample size (see Fig. 1-4). Significant sample sizes were defined as those with averaged t-values greater than the 97.5% quantile of the t-distribution.

#### Dynamics of α burstiness

We analyzed α burst dynamics using 25 s non-overlapping windows. Pairwise t-tests were performed to evaluate variations of the relative burst time across these windows. P-values were corrected for multiple comparisons using Holm’s method. To investigate the effect of this stability on the predictive power of the relative burst time, we computed Spearman’s correlation between the relative α burst time at each window and the relative duration estimated by the participant on the entire recording (rTE). Statistical significance was assessed with a one-sided test (rho > 0). Critical values for statistical significance were determined using tabulated references (Ramsey, 1989).

#### Stability of α markers before and after a timing task

A subset of participants from Exp. 2 (N = 21) also performed a resting-state EEG recording before performing a timing task, for a total of 90 s eyes open and 90 s eyes closed. To assess changes in spontaneous oscillatory dynamics before and after the task, a one-way repeated measures ANOVA was performed on the relative α burst. The first 90 s of the retrospective duration resting-state recording was used to match durations with the pre-recording resting-states. Post-hoc comparisons were performed using a pairwise t-test with Holm’s correction. Furthermore, to evaluate if changes in α affected channels in our cluster of interest, we ran a permutation t-test contrasting the average relative burst time during the eyes open recording before a task and after a task. The permutation procedure mirrored the spontaneous oscillation localizer method (see above), substituting integrated power with the relative burst time differences between conditions.

## Results

### Behavioral results

Participants retrospectively estimated the duration of a 4-minute resting-state EEG recording before (Exp. 1, N = 26) or after (Exp. 2, N = 45) performing a 90-minute timing task, or while seated in a virtual room (Exp. 3, N = 57). We computed the relative time estimates (rTE) as the ratio between the retrospective duration and the clock duration. A rTE of 1 signifies that participants correctly estimated the target duration, a rTE < 1 that they underestimated and a rTE > 1 that they overestimated (Fig. 1D).

On average and as predicted, participants in Exp. 1 significantly underestimated the duration of the recording (M = 0.80, SD = 0.24), as confirmed by a Wilcoxon signed-rank test (W = 24.00, p = 0.001).

Participants in Exp. 2 either performed a temporal adaptation task (N = 25) or an implicit timing task (N = 23) before the resting-state constituting the retrospective duration to be estimated. Before merging these two datasets, we confirmed that there were no behavioral differences between these two groups of participants (W = 226.00, p = 0.546; see Extended Data Fig. 1-2). In Exp. 2, like in Exp. 1, participants significantly underestimated the duration of the recording (M = 0.90, SD = 0.30; W = 251.50, p = 0.020; Fig. 1D). The observed underestimation of elapsed duration suggests that participants were unlikely to be paying attention to time during the resting-state recordings.

Contrary to Exp. 1 and Exp. 2, we found three outliers in the distribution of duration estimates (see Methods). No significant underestimations of elapsed duration in Exp. 3, when participants performed the resting-state recording in either virtual rooms (M = 0.94, SD = 0.33, W = 654, p = 0.171). Furthermore, we found no significant differences in the duration estimates between participants seated in the small VR (N = 29) and those seated in the large VR (N = 28) (W = 419.5, p = 0.835; Fig. 2C).

In addition to estimating durations, participants were also asked to judge their felt speed of the passage of time on a Likert Scale, ranging from “Very Slow” to “Very Fast”. We conducted an ordinal logistic regression with the PoTJ as the dependent variable and the experiment as the predictor (Extended-Data Fig. 1-4). The model has a significantly better fit than the null model (χ^2^(1) = 10.46, p = 0.005). At the predictor level, the study type was a significant predictor (Exp. 2: beta = −1.43, SE = 0.47, z = −3.07, p = 0.003), showing that participants in Exp. 2 rated their felt passage of time to be faster than participants in Exp. 1 and in Exp. 3 (Extended-Data Fig. 1-4). This observation will be important to account for the EEG observations in Exp. 2. The felt passage of time in Exp. 1 and Exp. 3 did not significantly differ (Extended-Data Fig. 1-4).

### EEG replication: α burstiness correlates with retrospective duration estimates

Using the cluster of parietal channels obtained with the α localizer, we averaged the relative α burst time over these channels to get one data point per participant in Exp. 1. We computed the linear model with α burst time as the main predictor of the rTE, which was statistically significant (R^2^ = 0.27, F(1, 24) = 8.86, p = 0.007). A significant positive relation between α burst time and rTE was found (β = 0.01, 95% CI [3.06e-03, 0.02], t(24) = 2.98, p = 0.007). Seeing that the correlational quantification of relative α burst time and rTE entails a single measure per participant, extreme inter-individual variability could disproportionately skew this relation. One participant was considered influential in this model (Cook’s Distance returned D = 0.25 with D > 4/N = 0.15) but the removal of this data point from the model did not affect the outcomes (β = 0.01, 95% CI [5.16e-03, 0.02], t(23) = 3.71, p = 0.001). We thus confirmed the relation between α burst time and rTE using Spearman’s correlation coefficient on the entire dataset (ρ(24) = 0.54, p = 0.005; Fig. 1E).

To assess the reliability of our observations, we asked how many participants would be required to conclude on the presence of an effect. The sensitivity analysis of the linear model showed that a minimum of 14 participants was necessary to detect the effect in Exp. 1 (Fig. 1-4). For this sample size, the t-threshold equated to 2.18, and the average observed t-value across the 100 repetitions was 2.23.

In sum, the observation of a significant positive correlation between relative α burst time and rTE observed in Exp. 1 replicated the original MEG findings(Azizi et al., 2023) and extended them to EEG. We then turned to Exp. 2, in which data were collected at the end of an experimental session.

### α burstiness does not predict retrospective duration after timing tasks

Prior to merging the two groups of participants from Exp. 2, we checked that the relative α burst times were comparable in the two groups. Using a Wilcoxon rank-sum test, we found no significant differences in relative α burst time between the two groups of participants (W = 278.00, p = 0.581). Data from all participants were thus merged into a single experimental group.

We investigated the relationship between relative α burst time and rTE in Exp. 2 using a similar approach as the one used in Exp. 1. The linear model with α burst time as the main predictor of the rTE was not significant (R^2^ = 0.002, F(1, 43) = 0.07, p = 0.788). Two participants were influential observations in Cook’s sense, and the removal of these two data points did not change the outcome of the model (R^2^ = 0.004, F(1, 41) = 0.18, p = 0.675). Overall, we found no significant correlation between rTE and relative α burst time in Exp. 2 (ρ(43) = −0.12, p = 0.420; Fig. 1F). We also performed the analysis separately for each group but found no significant correlations, whether participants took part in an explicit temporal adaptation task (ρ(21) = −0.18, p = 0.420) or in an implicit timing foreperiod task (ρ(20) = −0.11, p = 0.635) before the quiet wakefulness of the retrospective duration task.

In Exp. 3, 57 participants waited for 4 minutes in either a small or a large virtual room (Fig. 2A, balanced sample). As for Exp. 2, we checked that the relative α burst times were comparable in the two experimental conditions. First, we averaged the relative burst time over the channels of the parietal cluster (Fig. 2B), which was comparable to the one found in Exp. 1 and Exp. 2 (Fig. 1B) with a few more occipital sensors. Then, using an unpaired t-test, we found no significant differences in relative α burst time between the two groups of participants (t = 0.162, p = 0.872).

Additionally, we checked that the relative α burst times were comparable with Exp. 1, in which participants also performed the resting-state recording before any task. We found no significant differences between the two groups of participants (t = 0.96, p = 0.158).

Last, we investigated the relationship between relative α burst time using similar approaches as the one used in Exp. 1 and Exp. 2. The linear model with α burst time as the main predictor was not significant (R^2^ = 0.05, F(1, 55) = 2.74, p = 0.104). No participants were influential observations in Cook’s sense. Overall, we found no significant relation between rTE and the relative α burst time in Exp. 3 (ρ(55) = −0.19, p = 0.155; Fig. 2D).

### Specificity of α burstiness for the prediction of retrospective duration estimates

α burst dynamics may be a selective predictor of rTEs if no other part of the oscillatory spectrum predicts rTEs. To verify this, we performed the same correlation analysis as we did with α but this time, using slower (θ) or faster oscillations (β) known to present bursty dynamics (S. R. Jones, 2016; Little et al., 2019). Both θ and β were detected through a localizer approach, as was done for the α (see Methods).

θ bursts were located in frontal channels and θ burst time showed no correlation with the rTE in either experiment (Exp. 1 ρ(22) = 0.10, p = 0.636 in Fig. 3A; Exp. 2 ρ(42) = 0.01, p = 0.924 in Fig. 3C; Exp. 3 ρ(54) = −0.04, p = 0.783 in Fig. 3E).

**Figure 3.**
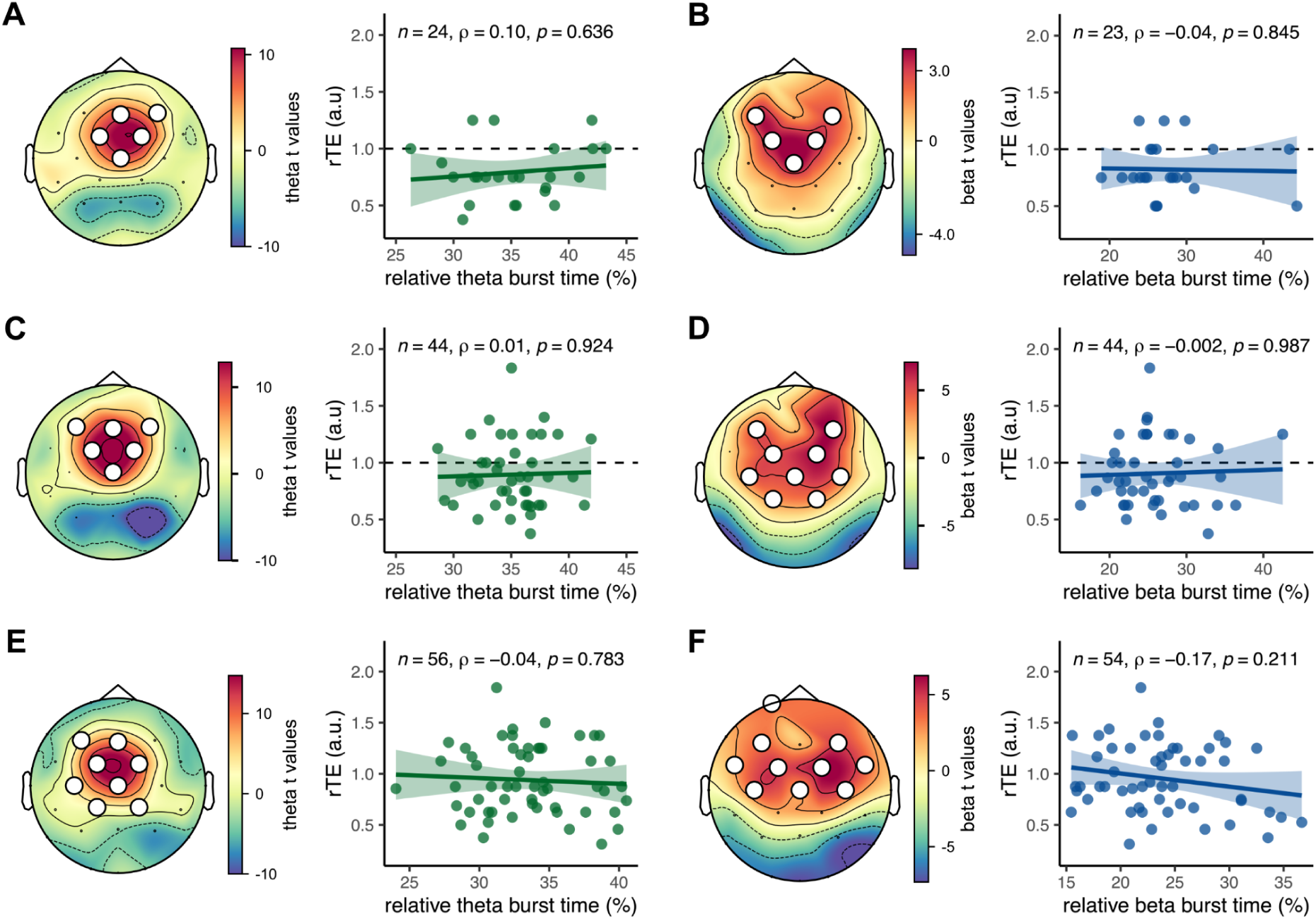
No correlation between relative θ or β burst time and rTEs. **(A)** *Left.* Localization of θ band activity using a permutation t-test on the z-scored integrated power in Exp. 1. *Right.* Spearman’s correlation between the relative θ burst time and the rTE in Exp. 1. The straight line is the regression line, and the shaded area is the 95% CI. The dotted line denotes the identity line between subjective and objective duration. **(B)** Same as (A) but for the β band in Exp. 1. **(C-D)** Same as (A-B) but in Exp. 2. **(E-F)** Same as (A-B) but in Exp. 3.

Similarly, β burst activity clustered in a larger set of frontal channels, and no significant correlations were found between relative β burst time and rTE (Exp. 1 ρ(21) = −0.04, p = 0.845 in Fig. 3B; Exp. 2 ρ(42) = −0.002, p = 0.987 in Fig. 3D; Exp. 3 ρ(52) = −0.17, p = −0.211 in Fig. 3F).

### The dynamics of α activity are stable during resting-state

We then investigated putative differences in electrophysiological data in the course of the experiment to understand why post-task α could not predict retrospective duration. Azizi et al.(2023) found that the first two minutes, and even the first 30 seconds, of the MEG signal were sufficient to predict participant retrospective duration estimates. This effect was explained by the stability of the α burst time over their resting-state recordings. Thus, we checked whether the α burst time recorded with EEG was stable in Exp. 1 and in Exp. 2, and compared them for eventual differences, which could account for the absence of replication in Exp. 2.

First, we tested whether relative α burst time changed over time (Fig. 4A). A pairwise t-test with Holm’s correction for multi-comparisons revealed that only the first window (0 to 25s) had a significantly lower burst time than all other windows in both Exp. 1 (t(25) =< −3.59, p < 0.006) and Exp. 2 (t(44) =< −3.91, p < 0.001). A Spearman’s correlation test between the relative α burst time in each window and the participants’ rTE on the whole recording revealed a significant positive correlation in Exp. 1 (0.34 ≤ ρ(24) ≤ 0.54; Fig. 4B, red line). All coefficients exceeded the critical value c = 0.331 corresponding to the one-sided test (ρ > 0; α = 0.05; N = 26; Ramsey (1989)). To the contrary, no correlations were found in Exp. 2 (−0.22 ≤ ρ(43) ≤ 0.03; Fig. 4B, pink line) seeing that all coefficients were inferior to the critical value c = 0.248 corresponding to the one-sided test (ρ > 0; α = 0.05 and N = 45). Last, and similar to the observation in (Azizi et al., 2023), the correlation coefficients in Exp. 1 slightly decreased over time, while remaining significant all along.

**Figure 4.**
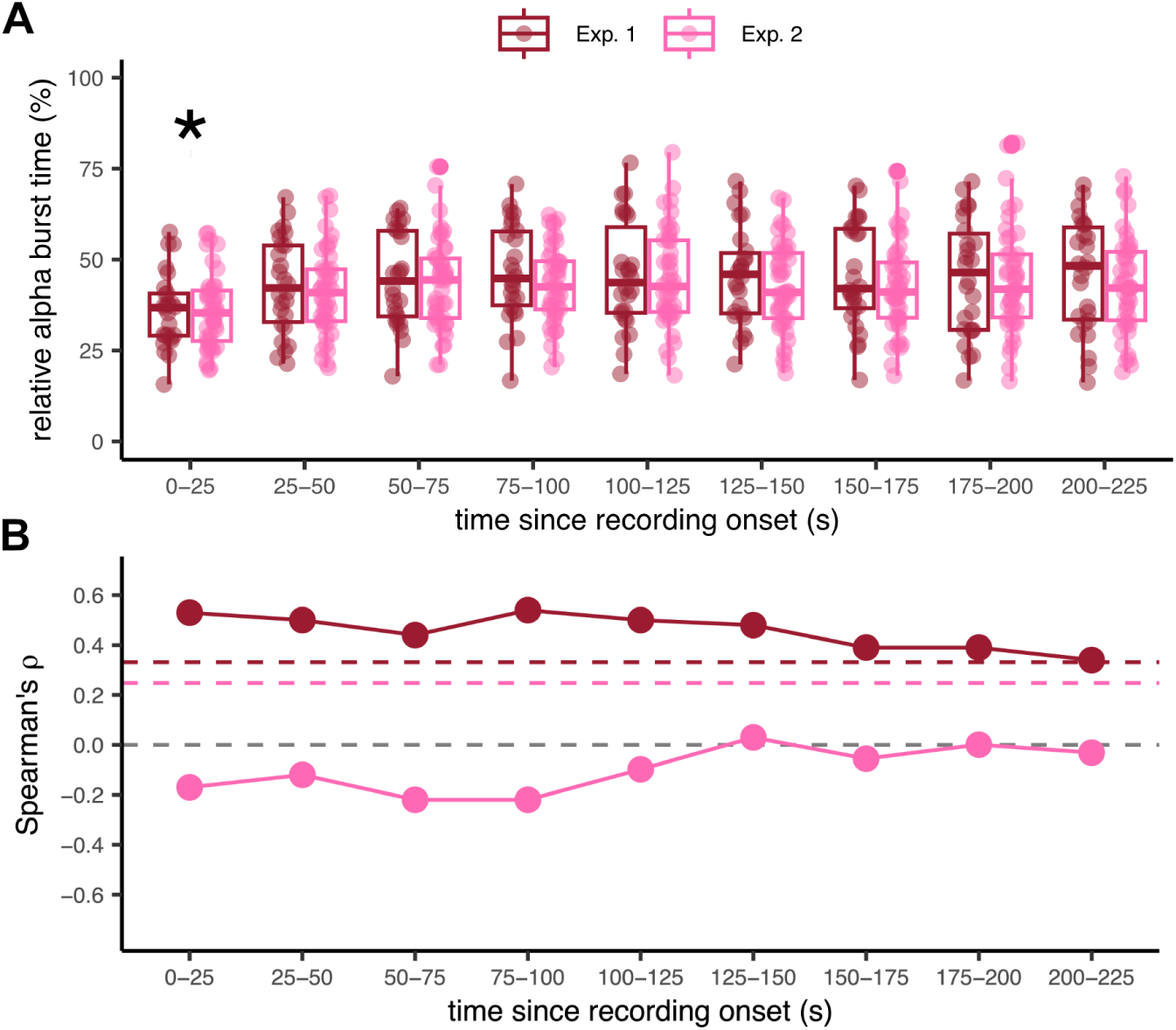
Correlation of binned α burst time and rTEs, using non-overlapping windows of 25 s (Exp. 1 and 2). **(A)** Stability of the relative α burst time over time by windows of 25 s for Exp. 1 (red) and Exp. 2 (pink). Each dot is a participant. According to a pairwise t-test with Holm’s correction, only the first 25s window differed significantly from all other windows in both experiments (*p < 0.002). **(B)** Spearman’s correlation coefficient between the relative α burst time for each window (as shown in (A)) and the rTE for Exp. 1 (red) and Exp. 2 (pink). The grey dotted line is the ρ = 0 line. For reference, the colored dotted line corresponds to the critical values of the one-sided tests (ρ > 0; α = 0.05) for N = 26 (Exp. 1, red line) and N = 45 (Exp. 2, pink line).

### Modulation of α activity over time and time-on-task effects

Time-on-task effects are systematic behavioral and neurophysiological changes that accompany the time spent on a given task. Recent work has described that time-on-task effects were associated with an increase in α power and a decrease in individual α peak frequency (iAPF) in the course of an experiment (Benwell et al., 2019). Herein, we investigated whether the lack of replication in Exp. 2 resulted from time-on-task effects in the α activity.

In Exp. 2, we also made 90s resting-state measurements at the beginning of the recording session, which allowed assessing intra-individual changes in the evolution of α throughout the experimental session. To directly test for the time-on-task effects and their possible associated changes in α dynamics, we quantified α power and iAPF in the resting states collected before and after the experimental session. The power of the α peak was significantly higher after the experimental session (M = 22.46, SD = 11.32) than it was before (M = 19.15, SD = 12.11), as assessed by a paired t-test (t(20) = 2.861, p = 0.010). No significant changes were found in the iAPF recorded before (M = 10.30, SD = 0.47) and after a timing task (M = 10.20, SD = 0.45; t(20) = −0.933, p = 0.362). Hence, we partially found time-on-task effects associated with α changes of power but not frequency.

Next, we asked whether the increased α power was caused by changes in relative burst time, seeing that two characteristics of α burst activity can cause an increase in spectral power: the average burst amplitude and the relative burst time (Azizi et al., 2023; Donoghue et al., 2022; Feingold et al., 2015; Jones, 2016). First, we quantified possible changes in burst amplitude before and after the experimental session. For this, we used a Wilcoxon signed-rank test for the contrast because the distribution of the burst amplitude recorded before the tasks did not follow a normal distribution (Shapiro’s test; W = 0.88, p = 0.011). Contrasting the average burst amplitude over the parietal cluster (Fig. 1B), we found that the burst amplitude was significantly larger after (M = 9.69 µV, SD = 3.92 µV) than before an experimental session (M = 9.04 µV, SD = 4.27 µV; W = 35.0, p = 0.004). Second, we compared the average relative burst time in the parietal cluster before (M = 40.34, SD = 10.64) and after (M = 41.53, SD = 10.97) the experimental session using a paired t-test. We found no significant differences (t(20) = −0.86, p = 0.400; Cohen’s d for paired samples of d = −0.19). However, when contrasting the relative α burst time before and after the experimental session, we found a significant cluster of three channels in the occipital area (Fig. 5A). None of these three channels overlapped with the original localizer cluster for α (Fig. 1B). The relative α burst time over this three-channel cluster did not correlate with the rTE either (ρ(20) = −0.19, p = 0.376; Fig. 5B). Taken altogether, these observations suggest that α activity is partially affected by time-on-task effects and that there are additional contributors to the generation of α activity recorded at the scalp level at the end of experimental sessions.

**Figure 5.**
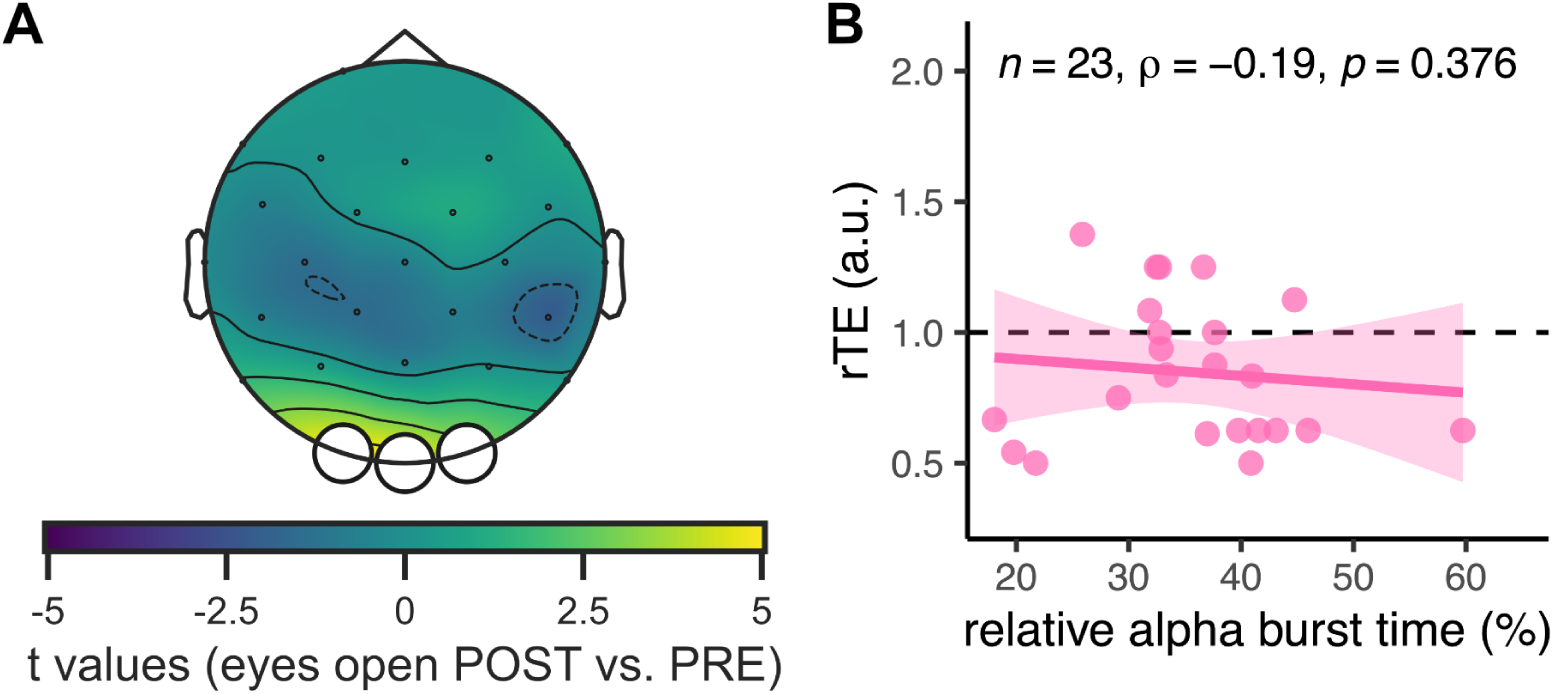
Additional contributors to α activity in Exp. 2 (N = 21). **(A)** Topographic map of t-values contrasting the relative α burst time in eyes-open resting states before and after an explicit timing task in a subset group of Exp. 2 (N = 21). A significant cluster (white dots) was identified, showing an increase in α relative burst time. **(B)** Spearman’s correlation between the relative α burst time averaged over the three channels of panel A and the rTE in the subset of Exp. 2 (N = 21). The straight line is the regression line, and the shaded area is the 95% CI. The dotted line denotes the identity line between subjective and objective duration.

## Discussion

In this series of EEG recordings, we replicate the original MEG observation reporting a positive linear correlation between resting-state α burst time and retrospective time estimation (Azizi et al., 2023). This replication suggests that the generators of α contributing to episodic timing can be captured non-invasively with both neuroimaging techniques. As in the initial report, this correlation was selective to α activity, and no such correlations were found for θ or β burst times during resting-state, despite their clear localization in the EEG spectrum and topographies. Intriguingly however, the replication only held when resting-state and retrospective timing estimates were collected before any tasks but disappeared when collected after participants performed timing tasks for ∼90 minutes. Additionally, we failed to replicate the correlation between α burst time and rTE when participants were seated in VR before any task. We discuss below the diverse reasons for replication and non-replication in this study.

### α dynamics recorded with EEG and MEG

One aim of the study was to replicate and generalize prior MEG observations to EEG. EEG remains a cost-effective technique and an invaluable tool for clinical applications and real-life mobile experimentation (Gramann et al., 2014; Stangl et al., 2023). Although MEG and EEG are both non-invasive and millisecond-resolved neuroimaging techniques, they differ in signal properties. MEG records the magnetic component of neural currents, which is less distorted by volume conduction through the dura and skull layers than the EEG signals (Baillet, 2017). This typically yields a higher signal-to-noise ratio in MEG, allowing experimental effects to be observed at the scale of an individual participant, whereas EEG tends to require population averaging. The α episodic marker investigated herein is an inter-individual correlational measure that could be highly sensitive to these differences in signal-to-noise ratio. With results of Exp. 1, we nevertheless replicate the original observations seen in MEG suggesting that the signal-to-noise of EEG is sufficient even with a reasonably low sample size (i.e., below 30 participants).

The sensitivity profiles between MEG and EEG also differ: EEG can theoretically detect sources in any orientation, whereas MEG is less sensitive to radial sources (Ahlfors et al., 2010). EEG tends to be more sensitive to neural generators located in gyri, and MEG to sulci. As the relation between α burst time and retrospective time holds despite these theoretical differences, this suggests that the neural sources contributing to the elicitation of spontaneous α bursts are either mostly radially oriented or largely distributed across cortex.

The sources of spontaneous α activity recorded with MEG and EEG are typically seen in occipital and parietal sensors, and source-localized in occipito-parietal regions (e.g. Azizi et al., 2023, Figures 1–2 Extended Data; Haegens et al., 2014; Salmelin & Hari, 1994). Still, this is a coarse approximation of the neural generators since such approach neglects the known modulations of posterior α activity through long-range and large-scale connectivity (Palva & Palva, 2007; Scheeringa et al., 2012). The neural generators of α activity have a long history (e.g. Adrian & Matthews, 1934): whether α generators are intrinsically cortical or driven by the thalamus is a debate initiated in the mid-twentieth century (Bremer, 1949; Kristiansen & Courtois, 1949). This question is still ongoing as both thalamocortical and intracortical generators of α activity appear to coexist. Corticothalamic α is well documented (Feige et al., 2005; Lőrincz et al., 2009; Vijayan & Kopell, 2012) and so is α activity in deep cortical layers (Buffalo et al., 2011; Buschman & Miller, 2007; X.-J. Wang, 2010). The propagation of α can also be cortico-thalamic (Halgren et al., 2019). With the current experimental design, we did not investigate the generative sources of α activity and future experiments will target a better understanding of neural sources contributing to the observed episodic timing effect, including the possible relation between α burst activity and α traveling waves (Das et al., 2022; Davis et al., 2021; Halgren et al., 2019). Nevertheless, it is likely that the α bursts characterized here reflect participants’ internal stream of thoughts during quiet wakefulness (Halgren et al., 2019; Sadaghiani & Kleinschmidt, 2016) or mind-wandering (Compton et al., 2019), and be elicited by as part of larger functional networks (Buckner & DiNicola, 2019; Raichle, 2015; Sadaghiani & Kleinschmidt, 2016).

### Absence of replication after task engagement: time-on-task effects and contextual changes

We replicated the relation between spontaneous α burst time and retrospective time estimate at the beginning (Exp. 1) but not at the end (Exp. 2) of an experimental session. The failure to replicate was not an outcome of trivial factors, which we thoroughly evaluated. For instance, the sample size exceeded the threshold recommended by our sensitivity analysis in the first experiment (Extended Data Fig. 1-4). Although we used two different EEG systems, their use was orthogonal to the conditions that were tested and, for instance, the data from half of participants tested in Exp. 2 were acquired with the same EEG settings as Exp. 1. Additionally, the localization of α burst activity in Exp. 2 was comparable to that observed in Exp. 1 (Fig. 1B) and in the original MEG study (Azizi et al., 2023). Thus, scalp-level α activity analyzed in Exp. 2 did not appear entirely different from the one analyzed in Exp. 1.

One major expected impact of testing participants at the end of an experimental session is mental fatigue, which has been associated with spectral changes in α but also θ activity (Boksem et al., 2005; Tran et al., 2020; Wascher et al., 2014). Time-on-task effects have also been reported in which α power increases, and α peak frequency decreases, in the course of an experimental task (Benwell et al., 2019). Seeing that the cerebral marker for episodic timing under investigation is α power, the increase of α power predicted by time-on-task should have led to an increase in retrospective time estimate or a possible disappearance of the linear relation between α and retrospective time estimation. As in the initial study (Azizi et al., 2023) and in Exp. 1, participants in Exp. 2 significantly underestimated the duration spent in resting-state. This supports the notion that participants did not pay attention to time during the recording, and that their behavioral trend was comparable to conditions before an experimental session. However, we did find partial evidence of time-on-tasks effects in EEG data with a significant increase in α power between the beginning and the end of a recording session. This α power increase predicted that participants would underestimate elapsed time less or perhaps overestimate it. This is not what we found. Interestingly, we did observe that the increase in α power was attributable to sensors that were not part of the original parietal α cluster, and that were instead more occipital. Thus, one possible explanation for the non-replication is that the neural generators contributing to the α recorded at the scalp may have changed in the course of the experimental recording. No anatomical MRI of participants was acquired, which prevented precise source reconstruction of the EEG data to disambiguate this aspect, but future work will address this.

One complementary explanation for the vanishing relation between α activity and retrospective time estimation at the end of an experimental session appeals to the contextual change hypothesis, which predicts that retrospective time estimation is influenced by the amount of contextual change experienced during the time interval (Block & Reed, 1978). Under this working hypothesis, the brain doesn’t track time *per se,* but instead reconstructs time based on the memory for changes or mental states that occurred during that lapse of time (Ornstein, 1969). Accordingly, duration estimates have often been reported to increase with the amount of contextual changes that have been experienced by participants (Block, 1978, 2014; Zakay & Block, 1997), high-arousal and emotionally salient events can also increase event segmentation thereby leading to longer remembered durations (Frederickx et al., 2013; J. Wang & Lapate, 2025; Zakay et al., 1994). At the beginning of an experimental session, the sequence of events is rather standard (welcoming to the lab, ethical approval, setting-up, explanations of the procedure) but more variability is expected by the end of an experimental session. One possibility is that the experimental context affected the storage and/or the retrieval of temporal information during the subsequent resting-state in a manner that we did not experimentally control for. The 90 minutes long experiments were segmented into shorter blocks and breaks, eliciting temporal clues and landmarks that could have influenced subsequent participants’ estimates, and also their resting-state dynamics. At the time participants received the question, contextual interference effects could impact the correlation between α episodic timing and retrospective duration estimate. Future studies will exploit this direction of research as the literature has shown that parametric changes during minute long experiences affect subsequent temporal memory (Brunec et al., 2020; Lositsky et al., 2016; Montchal et al., 2019). For instance, Brunec et al. (2020) showed that memory of duration increases with the number of turns that an individual took during spatial navigation. Similarly, (Lositsky et al., 2016) showed that the more event boundaries during an interval, the longer participants will estimate it to be.

Additionally, tasks tested in Exp. 2 involved timing, either explicitly or implicitly, which may have resulted in orienting participants’ attention to time. However, we think it is very unlikely because participants underestimated the retrospective duration as would be expected when not paying attention to time (Nicolaï et al., 2024; Polti et al., 2018). Recent findings also showed that participants who mind-wandered tend to underestimate the duration of events (Terhune et al., 2017). Still, the implication of the timing system for such a long recording period (90 minutes) may have influenced the dynamics of brain activity to the extent that resting-state dynamics during the episodic timing block, and the retrieval of stored information functionally dissociated. Although difficult to operationalize mechanistically, and perhaps unparsimonious, we cannot disregard this possibility, which requires empirical validation. This would require a particular experimental design in which the same participants are tested before and after taking part in timing tasks or in non-timing tasks.

A plausible explanation is that by the end of the experimental session, participants have developed temporal expectations for the duration of an experimental block. During the debrief of Exp. 2, 21.3% of participants effectively reported using the preceding experimental blocks or break times as a reference for their retrospective duration estimation. The temporal expectation response strategy would affect the decision stage so that the retrospective estimation of duration resulted from the comparison between stored duration memories of past blocks with the just elapsed one (Kaju & Maglio, 2022). This comparison strategy would account well for the observed behavioral outcomes: retrospective durations were underestimated not by lack of attention to time, but due to a shorter than expected experimental block. This strategy would also suggest that some participants may have paid attention to elapsing time during the recording, in which case the relation between α burst time and retrospective duration is predicted to vanish (Azizi et al., 2023). Prior studies (L. A. Jones, 2019; M. R. Jones & Boltz, 1989; Tanaka & Yotsumoto, 2017) predict that the passage of time would feel faster for participants in Exp. 2 than in Exp. 1 or Exp. 3 due to the existence of priors on the length of an experimental block. This is indeed what we observed (Extended-Data Fig. 1-4), suggesting that by the end of an experimental session, participants may have used such a comparison strategy. Implicit statistical learning and mind-wandering appear to be decoupled (Massar et al., 2020) so that the reported dissociation between α dynamics and retrospective duration observed here may result from the decisional aspect, not from the temporal dynamics of the mind-wandering (itself affected by time-on-task effects).

Last, adding on to time-on-task fatigue, context and the response strategy, the preceding task demands during the recording session could also influence the quality of subsequent resting-states and possibly alter the content of mind-wandering, which is another possibly exploitable line of research that we have not controlled for here.

Hence, for all practical purposes, it is important to better understand the context and the timing at which spontaneous α activity and retrospective time estimations are being recorded during an experimental setup. Retrospective timing tasks, as a single-trial and few-minutes long experiment, can easily be included in a series of experimental tasks (e.g. Chaumon et al., 2022). One clear recommendation at this stage would be to test retrospective timing tasks as early as possible in an experimental session to prevent fatigue, uncontrolled contextual effects on episodic timing or mind-wandering.

### Resting in a VR environment does not replicate

In Exp. 3, we wanted to test the generalizability of the episodic timing marker in VR and the possibility that the size of VR would impact time estimation and the marker. Neither the replication, nor the effect of VR size were found on retrospective duration or on its relation to α activity.

VR is being increasingly used in timing research (Jording et al., 2022; Lamprou-Kokolaki et al., 2024; Riemer et al., 2014; Tobin et al., 2010; Tobin & Grondin, 2009). One legitimate question is whether VR is comparable to real-life experiments (Bogon et al., 2024), an issue that has also been raised in spatial cognition (Taube et al., 2013). When estimating time intervals in the order of seconds, time in VR appears generally similar to real life (Bogon et al., 2024) but, interestingly, VR seems to alter the expectations about the duration of physical processes: for instance, how long a bottle takes to empty will differ in VR and in real-life, but not abstract time perception (Bogon et al., 2024). The realism of the VR environment can also affect time estimates, sometimes making them less accurate than in real life (Bogon et al., 2024; Mioni & Pazzaglia, 2023). Yet, emotional arousal and content impact subjective time perception suggesting that one’s experience of VR, rather than the environment itself, is what affects time perception (Mioni & Pazzaglia, 2023; van der Ham et al., 2019). This may imply large variability in how individuals relate to time in VR. Rutrecht et al. (2021) showed that in VR, the flow state or “loss of the sense of time” (Csikszentmihalyi, 2013) correlated with reduced time awareness and faster subjective time passage but did not correlate with the accuracy in retrospective duration estimation (Rutrecht et al., 2021). This corroborates that the subjective experience of the felt speed of the passage of time differs from the ability to accurately judge elapsed duration (Droit-Volet et al., 2017; Lamprou-Kokolaki et al., 2024; Wearden, 2015). As seen previously, if the two phenomenological outcomes and measures are distinct, their underlying mechanisms may not be fully unrelated.

The first major difference between Exp. 3 and other experiments is that participants did not significantly underestimate the duration spent in VR, whether in the small or in the large VR room. The lack of time underestimation in VR indicates that participants may have prospectively timed. Unlike in the other two experiments, participants were seated in a room where a lamp was placed in front of them. As the start of the experiment was indicated by the lighting of the lamp, this stimulation may have yielded prospective (implicit or explicit) timing and temporal expectations as to when the lamp would next be lit up. Alternatively, participants may also have experienced boredom, known to lengthen time estimation but also slow the felt passage of time (Droit-Volet & Meck, 2007; Watt, 1991), which is not what we found. Second, the size of the VR did not significantly impact the retrospective time estimation as would have been expected through other studies (DeLong, 1981; Riemer et al., 2014). Two differences can explain this incongruence: our study tested a different time scale and used a retrospective, not a prospective paradigm. Additionally, temporal distortions previously reported to be affected by the scale of the environment were realized in non-static environments. Hence, behavioral outcomes already predicted a possible lack of replication, seeing that an individual’s experience in a VR environment may already alter temporal phenomenology. Prospective timing was predicted to alleviate the relation between α and temporal estimation (Azizi et al., 2023).

It is perhaps noteworthy mentioning that the quality of EEG recordings while in VR was comparable to the other experiments, including the global statistics of α and relative α burst times. The topographic maps of α, θ and β frequency bands during quiet wakefulness were also consistent across experiments. Thus, the absence of relation between α bursts and retrospective time estimation is unlikely to be attributed to differences in signal quality.

### Conclusions

In three EEG experiments, we aimed to extend previous work demonstrating the involvement of oscillatory α bursts in episodic timing. We replicated the underestimation of the retrospective duration estimation of a resting-state and the positive linear relation between retrospective duration estimates and relative α burst time. However, we failed to find this relation when participants were tested at the end of an experimental session due to the contextual shaping of temporal expectations yielding different cognitive strategies in the estimation of retrospective duration. The relation did not generalize to VR, as participants may have likely oriented their attention to time in this environment. Overall, additional work is needed to assess the robustness of the link between spontaneous α activity and the tracking of episodic timing.

## Author contributions

VvW designed research; RB performed research and analyzed data; RB and VvW wrote the paper.

## Acknowledgements

We thank William Vallet (N=31), Christina Yi Jin and Anna Razafindrahaba (N=26), Clara Driaï (N=25), Camille Grasso (N=38), and Matthew Logie (N=27) for EEG data collection. We further thank Matthew Logie for his contribution to the VR setup and Leila Azizi for sharing code. We also thank the members of UNIACT at NeuroSpin for their help in recruiting participants.

## AI Disclaimer

During the preparation of this work, the authors used DeepL to help with the language as non-native speakers of English. After using this tool/service, all authors reviewed and edited the content as needed and take full responsibility for the content of the publication.

## Conflict of Interest

No

## Funding sources

The study was funded by the Project-ANR-18-CE22-0016 Wildtimes and the EXPERIENCE Project of the European Commission H2020 Framework Program Grant No. 101017727 to V.vW. Our lab is part of the DIM C-BRAINS, funded by the Conseil Régional d’Ile-de-France.

## Notes

### Competing Interest Statement

The authors have declared no competing interest.

