## Extended Data for "Spontaneous oscillatory activity in episodic timing: an EEG replication study and its limitations"

**Number of extended figures: 4**

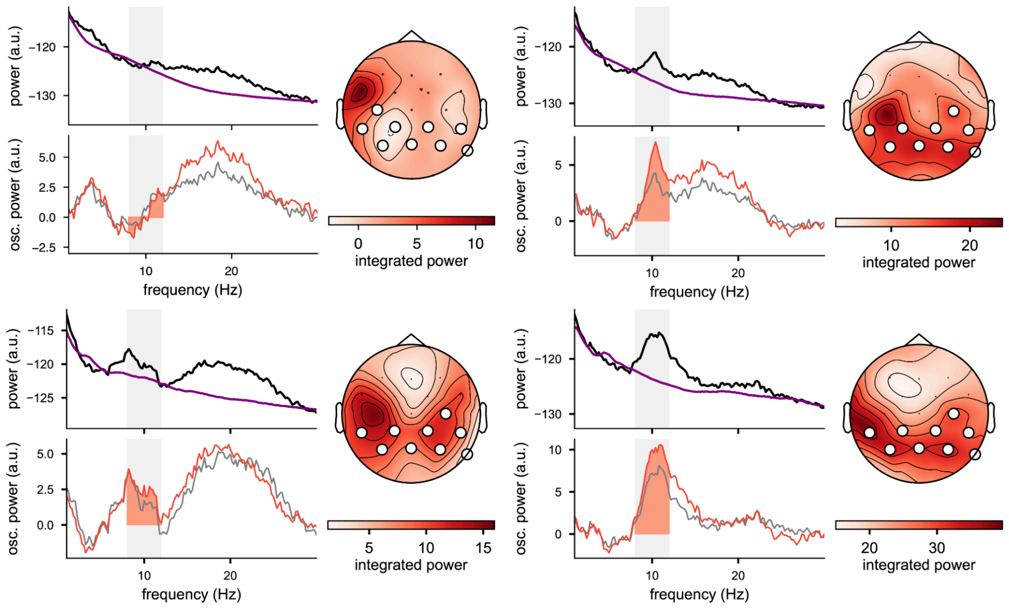


**Figure 1-1.** Topographic maps of excluded participants due to the lack of α activity in the cluster of channels localized by the cluster group.

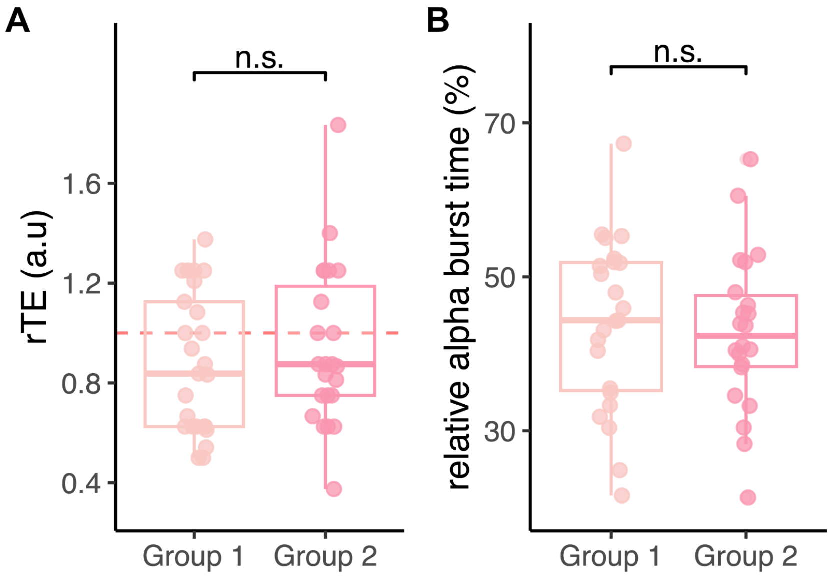


**Figure 1-2.** Exp. 2 data result from merging two homogeneous groups performing a resting-state EEG recording after a 90-minute timing task. **(A)** Behavioral distribution of Group 1 (after a temporal adaptation task) and Group 2 (after an implicit task). n.s. indicates a non-significant Wilcoxon test (W = 250.00, p = 0.443) **(B)** Bursts distribution of Group 1 (after a temporal adaptation task) and Group 2 (after an implicit task). ns indicates a non-significant Wilcoxon-test (W = 272.00, p = 0.677).

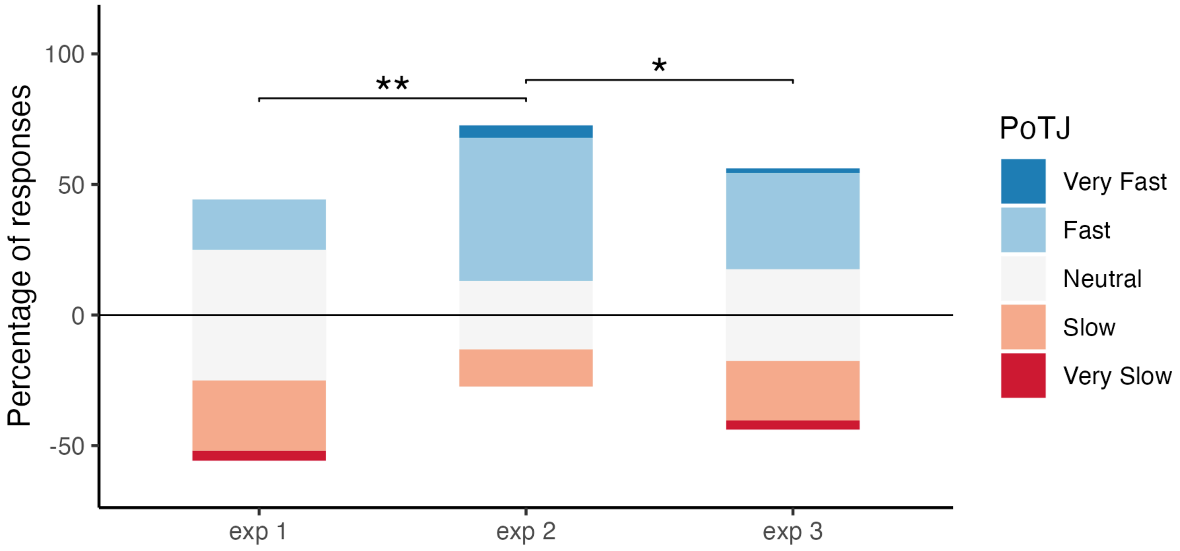


**Figure 1-3.** Felt passage of time judgments (PoTJ) were significantly faster in Exp. 2 than in Exp. 1 and in Exp. 3. Participants judged the experienced duration of the episodic time block on a Likert scale ranging from 1 (Very Slow, dark red) to 5 (Very Fast, dark blue). An ordinal logistic regression was computed with the PoTJ as the dependent variable and the experiment as the predictor. Significance was assessed through a likelihood ratio test against the null model (**p = 0.001; *p = 0.029).

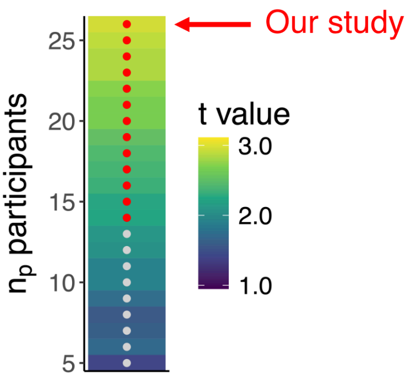


**Figure 1-4.** Sensitivity analysis of the relation between the retrospective time estimates (rTE) and the α bursts. Averaged t-value of the slope coefficient of the linear model for 100 repetitions of a sampling of $n_p$ participants. $n_p$ ranged from 5 to 26 (y-axis). Red dots indicate averaged t-values (over 100 repetitions) greater than the 97.5% quantile of the t distribution (with degrees of freedom of $n_p - 2$).
